# Disentangled Long-Read De Bruijn Graphs via Optical Maps

**DOI:** 10.1101/094235

**Authors:** Bahar Alipanahi, Leena Salmela, Simon J. Puglisi, Martin Muggli, Christina Boucher

## Abstract

Pacific Biosciences (PacBio), the main third generation sequencing technology can produce scalable, high-throughput, unprecedented sequencing results through long reads with uniform coverage. Although these long reads have been shown to increase the quality of draft genomes in repetitive regions, fundamental computational challenges remain in overcoming their high error rate and assembling them efficiently. In this paper we show that the de Bruijn graph built on the long reads can be efficiently and substantially disentangled using optical mapping data as auxiliary information. Fundamental to our approach is the use of the positional de Bruijn graph and a succinct data structure for constructing and traversing this graph. Our experimental results show that over 97.7% of directed cycles have been removed from the resulting positional de Bruijn graph as compared to its non-positional counterpart. Our results thus indicate that disentangling the de Bruijn graph using positional information is a promising direction for developing a simple and efficient assembly algorithm for long reads.

## 1 Introduction

Today, high-throughput DNA sequencing technology is central to every major (re)sequencing and *de novo* assembly project. Complex and long repetitive regions in genomes are a challenge for accurate assembly, especially for short-read sequencing technologies like Illumina, and this has driven a recent shift toward long-read sequencing technologies like Pacific Biosciences (PacBio). Long reads have already been successful in disambiguating the repetitive regions, leading to draft assemblies with fewer mis-assembled regions [25]. To date, however, long reads have a high error rate, which increases the complexity of assembly. For example, PacBio produces reads up to 50,000 bp in length, but with an insertion/deletion error rate of 15-20% [13].

Most assemblers targeting short read technologies use the Eulerian approach [12, 24]. In this assembly paradigm [12, 24] all unique *k*-mers (substrings of length *k*) are first extracted from the set of reads. A de Bruijn graph is then constructed with a vertex *v* for every (*k* – 1)-mer present in the set of reads, and an edge (*v*, *v*′) for every observed *k*-mer in the reads with (*k* – 1)-mer prefix *v* and (*k* – 1)-mer suffix *v*′. A contig corresponds to a non-branching path through this graph. SPAdes [1], ABySS [28], and Velvet [31] are examples of short read assemblers using the Eulerian approach. This approach is computationally efficient but does not easily adapt to reads with a high error rate. Moreover, applying it to long reads seems to discard the long range information in those reads.

With the above-mentioned caveats in mind, the first assemblers for long reads have adopted the *Overlap-Layout-Consensus (OLC)* approach. OLC first calculates the overlap between all pairs (or a subset of the pairs) of sequence reads and builds an overlap graph (in which there is an edge between pairs of reads having highest overlap). Similarly to the Eulerian approach, contigs then correspond to the non-branching paths through this graph. The computational bottleneck in OLC is the computation of (approximate) suffix-prefix overlaps between reads, which becomes computationally infeasible when the number of reads and the error rate grows.

Optical mapping is another technology that has been proposed for solving the repetitive regions in genomes. A genome-wide optical map contains the approximate genomic location of each restriction site corresponding to one or more restriction enzymes. Put another way, the optical map is the sequence of locations corresponding to all the occurrences of a short nucleotide sequence (3-5bp, say) in the genome. Optical maps span significantly larger genomic regions than long reads. For example, the typical region covered by a genome wide optical map is 300 Kbp [9], as opposed to the 15 kbp average length for a long read. The large length of the regions covered by a genome-wide optical map and recent increased commercial availability^1^, have lead to a rise in both data generation and tool development [30,17,15].

In this paper we consider Eulerian assembly applied to long reads in the presence of optical map data. In particular, we propose to use genome-wide optical maps to disentangle a de Bruijn graph constructed from long read data. In our approach we first correct sequencing errors in the long reads and then align the reads to the genome-wide optical map. This alignment information is then incorporated into the de Bruijn graph by constructing a *positional de Bruijn graph*, which is constructed from a set of *positional k-mers* (*k*-mer sequences with approximate positions associated with them) rather than *k*-mers alone. Since this variant of the de Bruijn graph effectively creates a separate *k*-mer for each unique occurrence of it in the genome, a space-efficient representation is vital for the graph to be constructed and used. We devise a space-efficient representation of the positional de Bruijn graph by augmenting a succinct BWT-based de Bruijn graph data structure [3]. We implement this method in a tool called Koota.

More specifically, our contributions are as follows: (1) a new Eulerian approach for long read assembly that is based on the positional de Bruijn graph; (2) a space-efficient representation of the positional de Bruijn graph, and (3) the first long read-optical map hybrid assembler. Our experimental results demonstrate that the positional information greatly reduces the complexity of the de Bruijn graph. In this paper, we study this complexity in terms of the number of cycles in the graph—which, using standard genome assembly terminology, are referred to as *bulges* (undirected cycles) and *whirls* (directed cycles). For example, when constructed for *E. coli* (K-12 substr MG 1655), the graph contains fewer than five whirls—a sharp reduction from 1,940, which is the number of whirls present in the traditional de Bruijn graph. Further, we show this gap between the number of whirls in the de Bruijn graph and its positional counterpart is substantially larger in yeast.

The results of Koota on *E. coli* and yeast are compared to those of ABruijn [16] and Canu [14], two leading long read assemblers. Koota was efficient with respect to both memory and time. It used the least memory to assemble *E. coli*—1.18 GB as compared to 3.7 GB and 2.7 GB for Canu and ABruijn, respectively—and required less than 5 GB of memory and approximately 12 hours to assemble yeast. This is in contrast to ABruijn, which required over 48 hours and 15 GB to assemble yeast. Further, we show that Koota achieved the lowest mismatch rate for both yeast and *E. coli,* and had a competitive genome fraction. This later statistic demonstrates that the fraction of the genome in the graph is not reduced by the removal of whirls and complexities within the graph. Thus these results show that disentangling the de Bruijn graph using positional information is a promising direction to develop an efficient and simple algorithm for long read assembly. Lastly, we note that Koota is freely available at https://github.com/baharpan/cosmo/tree/positional.

## 2 Background and Related Work

*Optical Mapping* Optical mapping is a technology that generates ordered, high-resolution, restriction maps of an entire genome. Optical maps are produced by immobilizing DNA molecules on a plate and applying a restriction enzyme on the molecules. Restriction enzyme will cleave the molecules at a specific DNA pattern *E*. The molecules are then imaged and the length of the fragments between restriction sites can be measured from the image. An optical map of a sequence is thus a sequence *R* = *r*_1_, *r*_2_,…, *r*_n_ where each *r_i_* is the length of the fragment between consecutive restriction sites.

Given a DNA sequence *X* and an enzyme recognizing the restriction site pattern *E* we can create an *in silico* digested optical map of it by mimicking how the enzyme cleaves the DNA molecule. Let *i*_1_, *i*_2_,…, *i_k_* be the occurrences of *E* in *X*. Then the *in silico* digested optical map of *X* is *M*(*X*|*E*) = *i*_2_ — *i*_1_, *i*_3_ — *i*_2_,…, *i_k_* — *i*_*k*−1_. For example if *X* =ACGAGACGGTTACGTG and *E* =ACG then the occurrences of *E* in *X* are 1, 6,12 and *M*(*X*|*E*) = 5, 6.

Since 2015 several methods for alignment of optical mapping data have become available, including OPTIMA [30], Maligner [17], and OMBlast [15]. Previously optical maps have been used for genome assembly in SOMA [19]. SOMA is a Eulerian assembler that uses both sequence data and optical mapping data. It builds the de Bruijn graph from short sequence reads and uses the optical map to eliminate or promote paths in the de Bruijn graph.

*Long-Read Assemblers* Canu [14], HGPA [7] and MHPA [2] are long read assemblers using the OLC approach. Canu is a fork of Celera assembler [18], which uses tf-idf weighted MinHash and a sparse assembly graph construction on its overlapping strategy. HGPA uses the Celera assembler [18] for the assembly and performs self-correction of continuous long reads sequences (CLR). MHPA [2], uses a probabilistic, locality-sensitive hashing for overlapping long reads that also works along with Celera assembler [18]. Lin *et al.* [16] present an Eulerian approach to assembling long reads. Their tool, called ABruijn, uses the A-Bruijn graph frame work, which is a different definition of de Bruijn graph. It constructs the A-Bruijn graph from a set of *k*-mers that are deemed sufficiently frequent; then it uses a path extension paradigm to derive genomic paths from short-read paths during traversal of the A-Bruijn graph; finally, errors in the draft genome are corrected using an *OLC* approach. Lin *et al.* [16] demonstrate that in order to correctly assemble long reads, only a small number of the reads are actually needed. They extract all unique *k*-mers from the (long) sequence reads then remove all *k*-mers that have frequency below or greater specific thresholds, the rationale being that those with low frequency are erroneous and those with high frequency originate from repetitive regions. Building the de Bruijn graph with this smaller set of filtered *k*-mers removes whirls and bulges in the resulting graph and simplifies the assembly process. The hybrid assembly using both short and long reads has also been considered for example by Pendleton et al. [22]. They combine long single-molecule and short high-throughput sequences to generate a hybrid genome assembly, which they then use to determine single nucleotide variants and structural variations.

*De Bruijn Graph Representations* Fundamental to our method is the succinct data structure for the positional de Bruijn graph. Although, there is a significant amount of work in constructing succinct de Bruijn graph representations- one of the first such approaches was introduced by Simpson et al. [27] as part of the development of the ABySS assembler-this is the first such representation for this de Bruijn graph variant. Nonetheless, we give a brief description of some of these works due to their inspiration and relevance to the work in this paper. Conway and Bromage [8] reduced space requirements from the method of Simpson et al. [27] by using a sparse bitvector (by Okanohara and Sadakane [20]) to represent the (*K*)-mers (the edges), and used rank and select operations to traverse it. Minia, by Chikhi and Rizk [6], uses a Bloom filter to store edges. They traverse the graph by generating all possible outgoing edges at each node and testing their membership in the Bloom filter. The representation that most closely reflects our work is BOSS graph representation of Bowe, Onodera, Sadakane and Shibuya [3] which is based on the Burrows-Wheeler transform [4]. Lastly, Chikhi et al. [5] implemented the de Bruijn graph using an FM-index and *minimizers*.

## 3 Methods

In this section we present our method for disentangling the de Bruijn graph using a genome-wide optical map. We start by describing how the sequencing errors in the reads are corrected and how the long reads are aligned to the genome-wide optical map. Then we define the positional de Bruijn graph and describe our method for constructing it. The section concludes with a description of how contigs are extracted from the positional de Bruijn graph.

### 3.1 Error Correction and Alignment of Long Reads

Before we can construct the positional de Bruijn graph, each long read must be localized on the genome-wide optical map. This involves three subprocesses: error correction, *in silico* digestion, and alignment.

We apply two rounds of error correction to the long reads: once before, and once after aligning them to the genome wide optical map. In both rounds we use the LoRMA long-read error correction method [26]. We note that LoRMA is purely long-read based, and makes no use of short reads. LoRMA proceeds in three phases. First a de Bruijn graph based approach is used for rough correction. The regions of the reads deemed to be unrecoverable by the de Bruijn graph based method are then cut out. This process trims and splits the reads. In the final phase multiple alignments are formed between similar reads. In this work we used the intermediate reads after de Bruijn graph based correction to avoid splitting the reads.

After this first round of error correction, we create for each read *r* an *in silico* digested optical map *M*(*r*|*E*) where *E* is the restriction site pattern(s) recognized by the restriction enzyme(s) used to build the genome-wide optical map. We then align these *in silico* digested reads to the genome-wide optical map using the method by Valouev et al. [29]. Of the alignments returned by that method, we retain only those for which at least 40% of fragments align (this threshold was found experimentally, and reduces the number of clearly erroneous alignments reported by Valouev et al.’s software). A second round of error-correction is then applied to this subset using LoRMA. We saw superior results with the *E. coli* data (see Section 4) when we error corrected a second time, but results were not substantially improved by a third round of error correction. After these steps we have a set of error corrected reads and for each of these reads we have an approximate genomic position based on the alignment to the genome wide optical map. This set of aligned reads and their genomic positions will then be used to build the positional de Bruijn graph.

### 3.2 Succinct Positional de Bruijn Graph

If a read *r* = [*r*_1_…*r*_*n*_] of length *n* is aligned to optical map at position *i*, we extract *n* – *k* + 1 *positional *k*-mers* from *r*: ([*r*_1_…*r*_*k*_], *i*), ([*r*_2_…*r*_*k*+1_], *i* + 1),…, ([*r*_*n*–*k*+1_…*r*_*n*_], *i* + *n* – *k*). Next, we refer to the *multiplicity* of a positional *k*-mer occurring at position *p* as the number of the occurrences of that *k*-mer at *p*. We emphasize that different reads may give rise to the same *k*-mer with different inferred positions,which allows us to disambiguate *k*-mers at different positions. We further note that the alignment to the optical map also gives the orientation of each read and thus also the orientation of each *k*-mer. Therefore, unlike in a de Bruijn graph without positional information, there is no need to merge a *k*-mer and its reverse complement which simplifies the construction and processing of the graph.

The positional de Bruijn graph *G_k,Δ_* is defined for a multi-set of positional *k*-mers and parameter *Δ*, and constructed in a similar manner to the traditional de Bruijn graph using an A-Bruijn graph framework from [23]. Given a *k*-mer *s_k_*, let prefix(*s_k_*) be the first *k* — 1 nucleotides of *s_k_*, and suffix(*s_k_*) be the last *k* – 1 nucleotides of *s_k_*. Each positional *k*-mer (*s_k_*,*p*) in the input multi-set corresponds to a directed edge in the graph between two positional (*k* – 1)-mers, (prefix(*s_k_*),*p*) and (suffix(*s_k_*),*p* + 1). Due to indels in the reads, it is possible that different positional (*k* – 1)-mers actually correspond to the same original location in the underlying genome. Therefore after all edges are formed, the graph undergoes a gluing operation, where positional (*k* – 1)-mers are glued together as follows. We group together positional (*k* – 1)-mers having the same (*k* – 1)-mer sequence and positions within Δ of each other. Such a group of m positional (*k* – 1)-mers is then replaced with a single positional (*k* – 1)-mer having a position equal to the average position of the group. An associated multiplicity is also stored.

Disambiguating identical *k*-mers (with positional information) should lead to a simpler graph, but an overall increase in space usage is likely because, for example, the positional de Bruijn graph will have more nodes than the plain graph, so care must be taken with graph representation. We have implemented a space-efficient data structure for storing and traversing the positional de Bruijn graph that is based on the BOSS de Bruijn graph representation of Bowe, Onodera, Sadakane, and Shibuya [3]. We begin by briefly defining the BOSS construction of the de Bruijn graph and then demonstrate how this structure can be extended to allow positional information to be stored. The first step of constructing this graph *G* for a given set of *k*-mers is to add dummy *k*-mers (edges) to ensure that there exists an edge *k*-mer starting with first *k* – 1 symbols of another edge’s last *k* – 1 symbols. These dummy edges ensure that each edge in *G* has an incoming node. After this small perturbation of the data, a list of all edges sorted into right-to-left lexicographic order of their last *k* – 1 symbols (with ties broken by the first character) is constructed. We denote this list as F, and refer to its ordering as *co-lexicographic ordering.* Next, we define L to be the list of edges sorted co-lexicographically by their starting nodes with ties broken co-lexicographically by their ending nodes. Hence, two edges with same label have the same relative order in both lists; otherwise, their relative order in F is the same as their labels lexicographic order. The sequence of edge labels (*k*-mers) sorted by their order in list L is called the *edge-BWT* (EBWT). Now, let BF be a bit vector in which every 1 indicates the last incoming edge of each node in L, and let BL be another bit vector with every 1 showing the position of the last outgoing edge of each node in L. Given a character *c* and a node *v* with co-lexicographic rank rank(*c*), we can determine the set of *v*’s outgoing edges using BL and then search the EBWT(*G*) for the position of edge *e* with label *c*. Using BF we can find the co-lexicographical rank of *e*’s outgoing edge.

We augment the BOSS representation with extra information per edge to obtain a positional de Bruijn graph. In particular, we build and store a vector *V* of integer vectors (containing positions associated with each node). Integers in each vector are stored bit packed, using the SDSL library [10], which also provides us fast random access to individual positions. *V* is indexed by *k*-mer lexicographic rank, so that *V*[*i*] is the set (vector) of positions where the *i*th lexicographically ranked *k*-mer in the input occurs. Rank operations on EBWT allow us to easily map from positions in EBWT (edges in the de Bruijn graph) to associated sets of positions in *V*.

Figure 1 illustrates a small example of the positional de Bruijn graph representation built for a set of 4-mers and *Δ*=4.

**Fig. 1:**
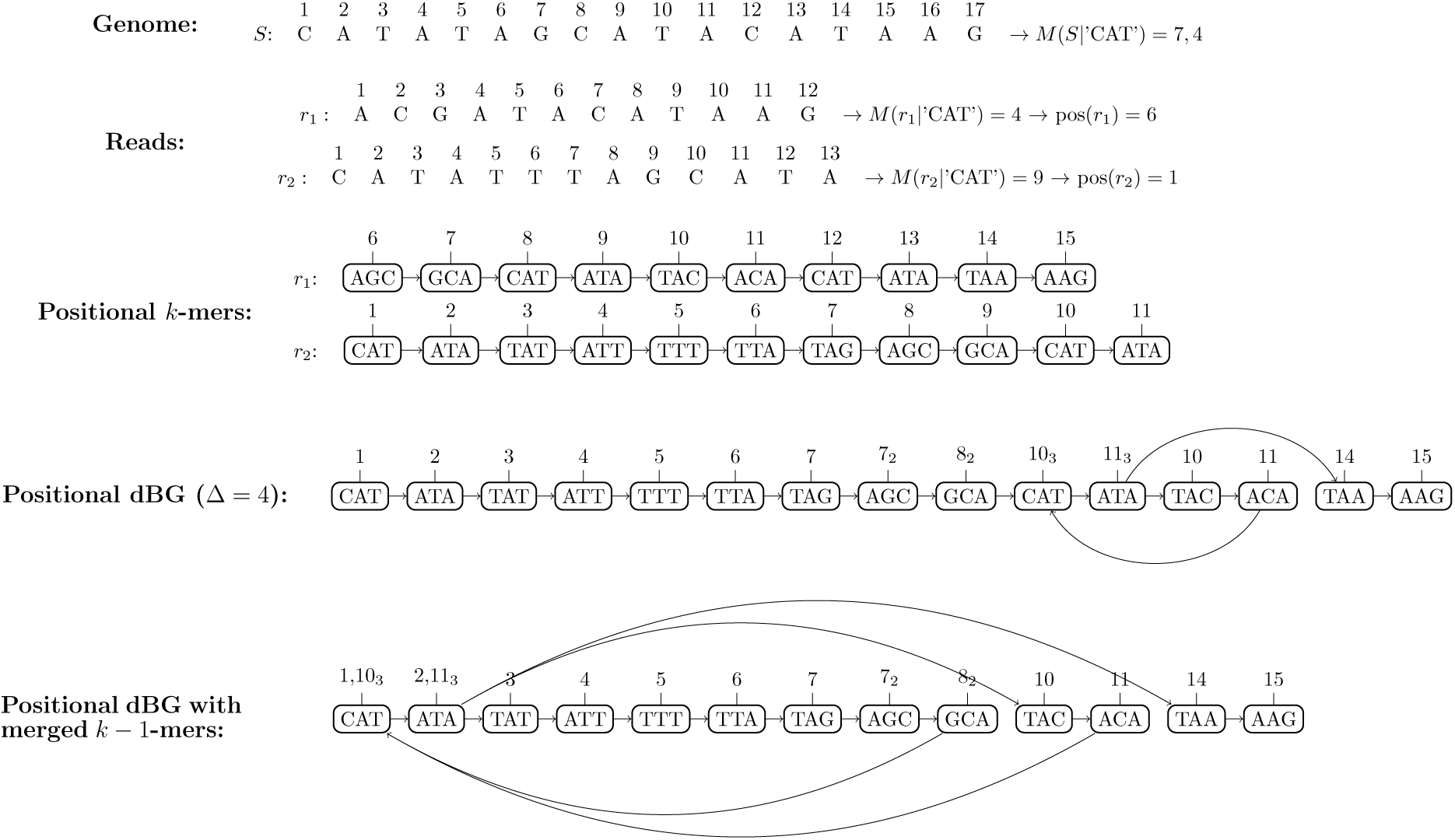
Positional de Bruijn graph *G*_4,4_ constructed from the reads *r*_1_ and *r*_2_ and the genome wide optical map *M*(*S*|’CAT’). Top of the figure shows the genome sequence *S* and the corresponding genome wide optical map. Note that *S* is not available to the method but it is shown here for clarity. Our method then *in silico* digests the error corrected reads *r*_1_ and *r*_2_ producing optical maps *M*(*r*_1_|’CAT’) and *M*(*r*_2_|’CAT’). They are aligned against the genome wide optical map yielding the positions 6 and 1 for the reads *r*_1_ and *r*_2_, respectively. The positional (*k* – 1)-mers obtained from the reads and their positions are shown in the middle. The positional de Bruijn graph is then constructed by gluing together (*k* – 1)-mers whose positions are within Δ = 4 of each other. The positions of the (*k* – 1)-mers are shown above the nodes and the multiplicity of the positional (*k* – 1)-mer is shown as a subscript if it is greater than one. Note that the positional (*k* – 1)-mers (AGC, 6) and (AGC, 8) are correctly glued together as they originate from the same genomic positions but are derived from different reads. On the other hand we see that although the positional (*k* – 1)-mers (CAT, 8), (CAT, 12), and (CAT, 10) do not all originate from the same genomic position, they are all glued together creating a small whirl in the graph. Bottom of the figure shows the positional de Bruijn graph where all positional (*k* – 1)-mers with the same (*k* – 1)-mer are merged to a single node with a list of positions and their multiplicities.

### 3.3 Construction of the Positional de Bruijn Graph

In our implementation, we first count the *k*-mers and calculate their associated positions (using the positional information of each read that comes from its alignment to the optical map). After clustering the positional *k*-mers as described in the previous section, we write the lexicographically sorted *k*-mers and the associated positions in separate files (both ordered by *k*-mer). Each *k*-mer is indexed by its lexicographic rank (lexrank). We build and store a vector of position sets *V*, in which, *V* [*i*] is the set of positions at which the *k*-mer with lexrank *i* occurs in the genome. Then we construct the BOSS representation for the *k*-mers such that instead of co-lexicographically sorting *k*-mers only, we sort (*k*-mer, lexrank) pairs.

To construct the F table, for each *k*-mer, (*k*-mer, lexrank) pairs will be sorted by the first *k* – 1 symbols (the source node of the edge). Similarly, to construct the L table, we also sort each *k*-mer (without its row number) by the last *k* – 1 symbols (the next node of the edge). At this step we calculate F \ L (comparing only the (*k* – 1)-length prefixes and suffixes respectively), and L\F to find the nodes that require incoming dummy edges and outgoing dummy edges, respectively. We will sort the set of incoming dummy edges by their first *k* – 1 symbols. We call this table *D*. The set of outgoing edges does not require sorting. Eventually we merge *D* with F and L \ F. During the merge we push the index of each resulting edge to a vector. Afterward, while traversing the *j*th edge (*k*-mer) in the graph, the *k*-mer’s index allows us to map to the *j*th element of the index vector, providing us access to the appropriate part of *V* containing the set of its associated positions of the *k*-mer. Sorting the *D* and F (arrays of *k*-mers) is the computational bottleneck in construction, and overall construction of the data structure takes *O*(*k*(|F|(log |F|)) time.

### 3.4 Graph Traversal and Contig Recovery

A contig in the positional de Bruijn graph is a non-branching path and thus to recover the contigs it is sufficient to enumerate all non-branching paths in the graph. The following procedure is repeated until all positional *k*-mers in the graph have been visited. We start by picking an unvisited positional *k*-mer (*s_k_*,*p*) and mark it as visited. We then traverse the graph both forward and backward starting from (*s_k_*,*p*). Let us consider the forward traversal. In our representation we need to retrieve all ouneighbors of the *k*-mer *s_k_*. We then filter the position lists of the outneighbors to find all positional *k*-mers (*s*′_*k*_,*p*′) such that *p*′ is within *Δ* of *p*. We say that these positional *k*-mers are consecutive to (*s_k_*,*p*). If a consecutive positional *k*-mer is marked visited or if there are no or several consecutive positional *k*-mers, we have reached the end of a non-branching path and stop our traversal. If there is exactly one consecutive positional *k*-mer, we mark that *k*-mer visited and continue the traversal from that *k*-mer. After the forward traversal finishes, we will traverse backward from the initial *k*-mer which proceeds analogously to the forward traversal.

## 4 Results

### 4.1 Datasets

We simulated 92,818 number of PacBio reads from the reference genome of *E. coli* K-12 substr. MG 1655 with model-based simulation of PBSIM [21] using the following parameters: mean accuracy of 85%, average read length of 10,000, and minimum read length of 1,000, and average coverage of 200x. According to observed distributions of real PacBio read length, the model-based method simulates PacBio reads with a log-normal length distribution. The average accuracy over the length of each read is taken from a normal distribution. We simulated an optical map using the reference genome for *E. coli* (str. K-12 substr. MG1655) and the enzymes XhoI, NheI and EagI since there is no publicly available one for this genome. The simulation was done by finding the locations of each restriction site in the reference genome and then *in silico* digesting at those locations.

Our second dataset consists of 220,336 sequence reads from *Saccharomyces cerevisiae* str. W303 (yeast) using data generated using PacBio RS II System and P4-C2 chemistry. The reads are available for public download from PacBio DevNet^2^. The average read length is 6,349 bp, with the minimum and maximum read length being 500 bp and 30,164 bp, respectively. Given there is no publicly avaliable optical map for yeast, one was simulated the *Saccharomyces cerevisiae* str. W303 and enzymes XhoI, NheI and EagI.

### 4.2 The Effect of Filtering and Error Correction

As previously mentioned, in order to use long reads in the construction of the positional de Bruijn graph, they need to be aligned to the genome-wide optical map. Error correction was used to maximize the number of reads that aligned to the optical map and thus, could be used for assembly. Prior to error correction, only 30% of the simulated *E. coli* reads aligned to the optical map, whereas 57% of them aligned to the optical map after error correction. Of these 57% of reads, 45% of them had an alignment where at least 40% of the fragments aligned. This increase in the aligned reads reflects the increase in the overall quality of the reads. The distribution of the frequency of *k*-mers changed dramatically with both the first and second error correction. This is illustrated in Figure 2. Prior to the first error correction a large portion of the reads had either very high frequency or very low frequency. We note that both of these sets of reads would be filtered by ABruijn. After the first error correction, alignment, and second error correction, the distribution of the *k*-mer frequency was much more uniform, with the majority of the *k*-mers having frequency between 20 and 90. Thus, as can be seen, the majority of these *k*-mers can be more effectively used for the assembly process by disambiguating them.

**Fig. 2:**
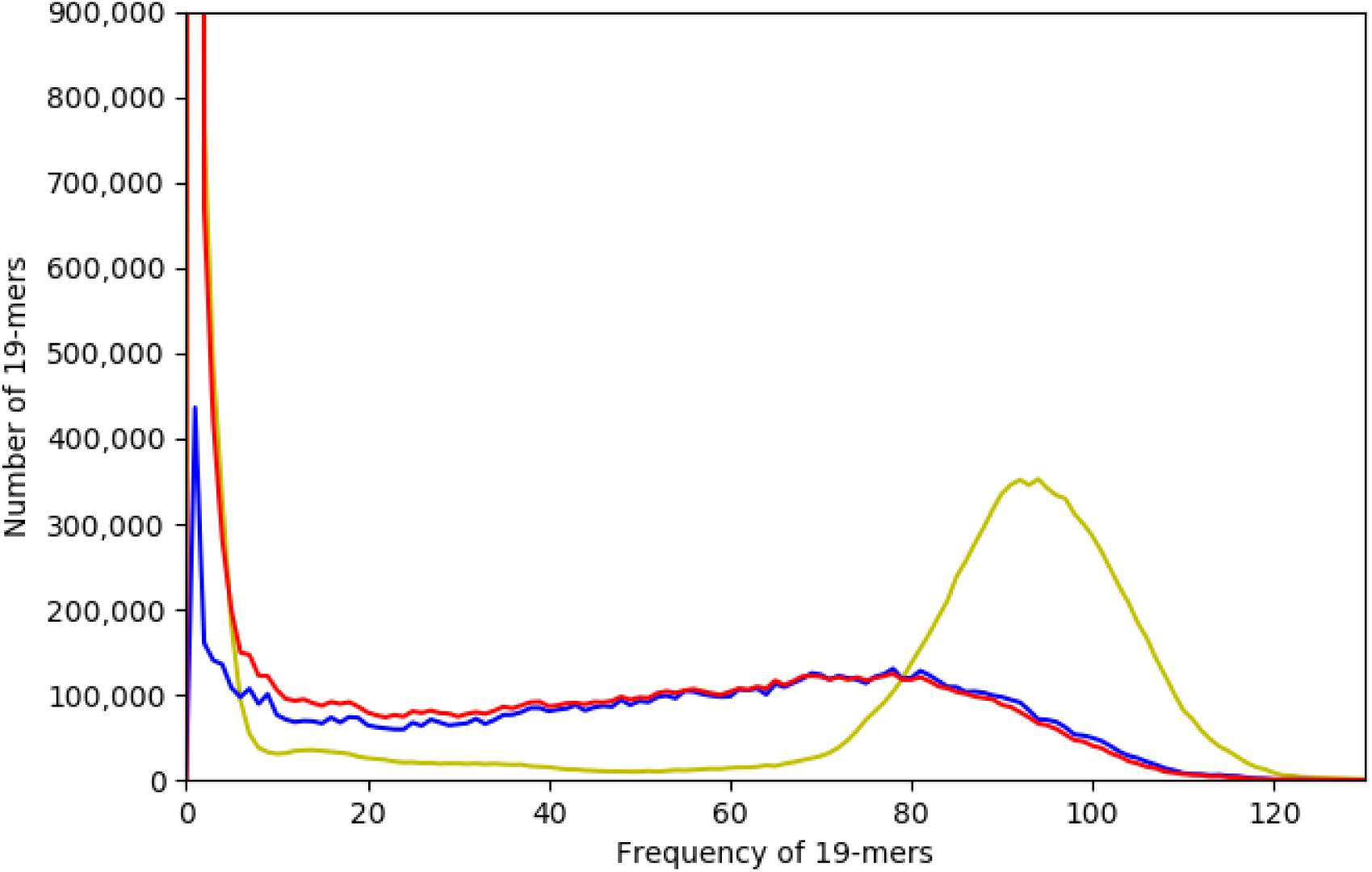
An illustration of the effect of error correction on number of aligned reads and 19-mer frequency. The histogram in yellow illustrates the frequency of all unique 19-mers in the initial corrected data. The histogram in red illustrates the frequency of all unique 19-mers that occur in reads that underwent one error correction and aligned to the genome-wide optical map. Lastly, the histogram in blue illustrates the frequency of all 19-mers that underwent error correction twice (once prior to alignment and once afterward) and aligned to the genome-wide optical map.

### 4.3 Comparison Between Assemblies

In addition to evaluating performance of Koota with respect to memory and time, we also analyzed the ability of Canu, ABruijn, Koota and to accurately assemble both datasets. All assemblers were ran with their default parameters, on the filtered, error corrected data and using *k* = 19. Koota used *Δ* = 500. All statistics were computed by QUAST in default mode [11]. The results demonstrate that Koota achieves the best *E. coli* assembly with respect to both genome fraction and rate of mismatches. Although the Canu and Koota *E. coli* assembly had similar genome fractions—93.25% and 94.23%, respectively— ABruijn had a much lower genome fraction (62.94%)—and Canu had a significantly higher number of mismatches than both ABruijn and Koota. The contigs constructed by Koota had a mismatch rate of 1.71 per 100 kbp. ABruijn and Canu had a mismatch rate of 1.16 and 2.89 per 100 kbp, respectively.

From 220,336 long read from yeast, 95,289 (approximately 43%) of them aligned to optical map, were error corrected a second time, and subsequently used for assembly. All the assemblies produced by Koota and ABruijn had similar genome fractions—92% and 93.5%, respectively; however, ABruijn had a substantially higher mismatch rate (90.47 mismatches per 100 kbp) than Koota (14.52 mismatches per 100 kbp). The Canu assembly had a moderately higher genome fraction (95%) in comparison to ABruijn and Koota but also a higher mismatch rate (22.9 mismatches per 100 kbp) in comparison to Koota.

### 4.4 Time and Memory Usage

We compared the resource usage of Koota with ABruijn and Canu on the two datasets, in particular peak memory usage, which was measured as the maximum resident set size, and run time, measured as the user process time. All experiments were performed on a 2 Intel(R) Xeon(R) CPU E5-2650 v2 @ 2.60 GHz server with 512GB of RAM, and both resident set size and user process time were reported by the operating system. Again, Canu, ABruijn, and Koota were applied to long reads that had undergone error correction and filtering. Table 1 shows the memory and time usage of the three different assemblers on both the *E. coli* and yeast datasets. The assembly time of Canu was moderately less than Koota. Canu required 7 minutes and 35 seconds to assemble *E. coli* and 3 hours and 35 minutes to assemble yeast; whereas, Koota required 32 minutes and 20 seconds to assemble *E. coli* and 12 hours and 4 minutes to assemble yeast. Both Canu and Koota also used less than 5 GB of memory to assemble both yeast and *E.coli.* Lastly, as can be seen in Table 1, ABruijn required more time and memory to produce assemblies for both *E.coli* and yeast.

**Table 1:**
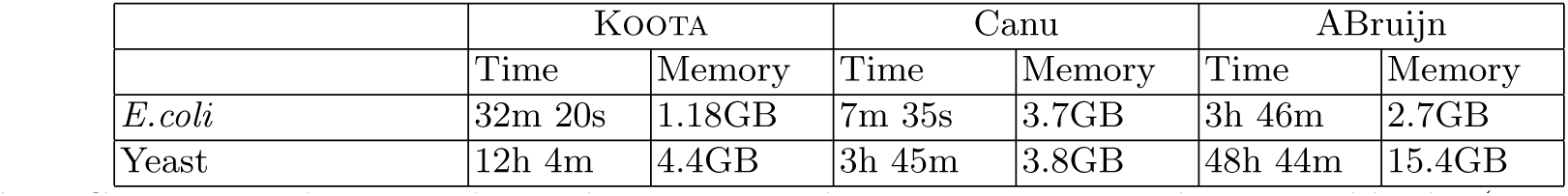
Comparison between the peak memory and time usage required to assemble the (error corrected and aligned) *E. coli* reads using Koota, ABruijn, and Canu. The reads were simulated using PBSIM [21] and *E. coli* K-12 substr. MG 1655 reference genome. k = 19 was used for all assemblers. The peak memory is given in megabytes (MB) or gigabytes (GB). The running time is reported in seconds (s), minutes (m), and hours (h).

|  | KOOTA |  | Canu |  | ABruijn |  |
| --- | --- | --- | --- | --- | --- | --- |
|  | Time | Memory | Time | Memory | Time | Memory |
| *E. coli* | 32m 20s | 1.18GB | 7m 35s | 3.7GB | 3h 46m | 2.7GB |
| Yeast | 12h 4m | 4.4GB | 3h 45m | 3.8GB | 48h 44m | 15.4GB |

## 5 Discussion and Conclusions

Development of a production quality assembler requires sophisticated traversal algorithms, the implementation of which is well beyond the scope of this paper. Our aim in developing Koota is to demonstrate that incorporating the approximate positions of the *k*-mers into the de Bruijn graph construction can greatly reduce the complexity of the resulting graph. Furthermore, using space-efficient encodings, the positional information can be added without a dramatic increase in memory requirements.

Koota required the least space to assemble the simulated *E. coli* reads; 1.18 GB in comparison to the 3.7 GB required by Canu and the 2.7 GB required by ABruijn. Koota also had the highest genome fraction of the methods tested, and a low mismatch rate. Taken together these statistics show that we have not discarded a significant portion of the genome, making accurate assembly possible. For completeness we report that Koota’s N50 scores are currently low (2,301 vs. 126,754 for Canu on the *E. coli* dataset), however this belies the absence of a sophisticated traversal algorithm to effectively deal with branches in the graph, and to resolve the remaining whirls and bulges. We reemphasize that our goal was not to compete with state-of-the-art assemblers, but instead to demonstrate how positional information can simplify the de Bruijn graph, in the context of long reads.

Indeed, the real influence of the optical map is its ability to disentangle the de Bruijn graph by assigning approximate positions to each of the long reads (and so the *k*-mers), and the addition of positions to the graph greatly reduces the number of whirls and bulges. To further illustrate this point, we constructed both the de Bruijn graph and positional de Bruijn graph on the set of filtered, error-corrected simulated *E. coli* long reads (see Subsection 4.1 for a full description of this data) with *k* = 19. We then counted the number of whirls where each *k*-mer or positional *k*-mer has multiplicity greater than one in each graph. The number of such whirls in the de Bruijn graph and positional de Bruijn graph was 1,940 and 3, respectively. Similarly, for the yeast genome the de Bruijn graph had 13,223 whirls, whereas the positional de Bruijn graph had only 302 whirls.

**Fig. 3:**
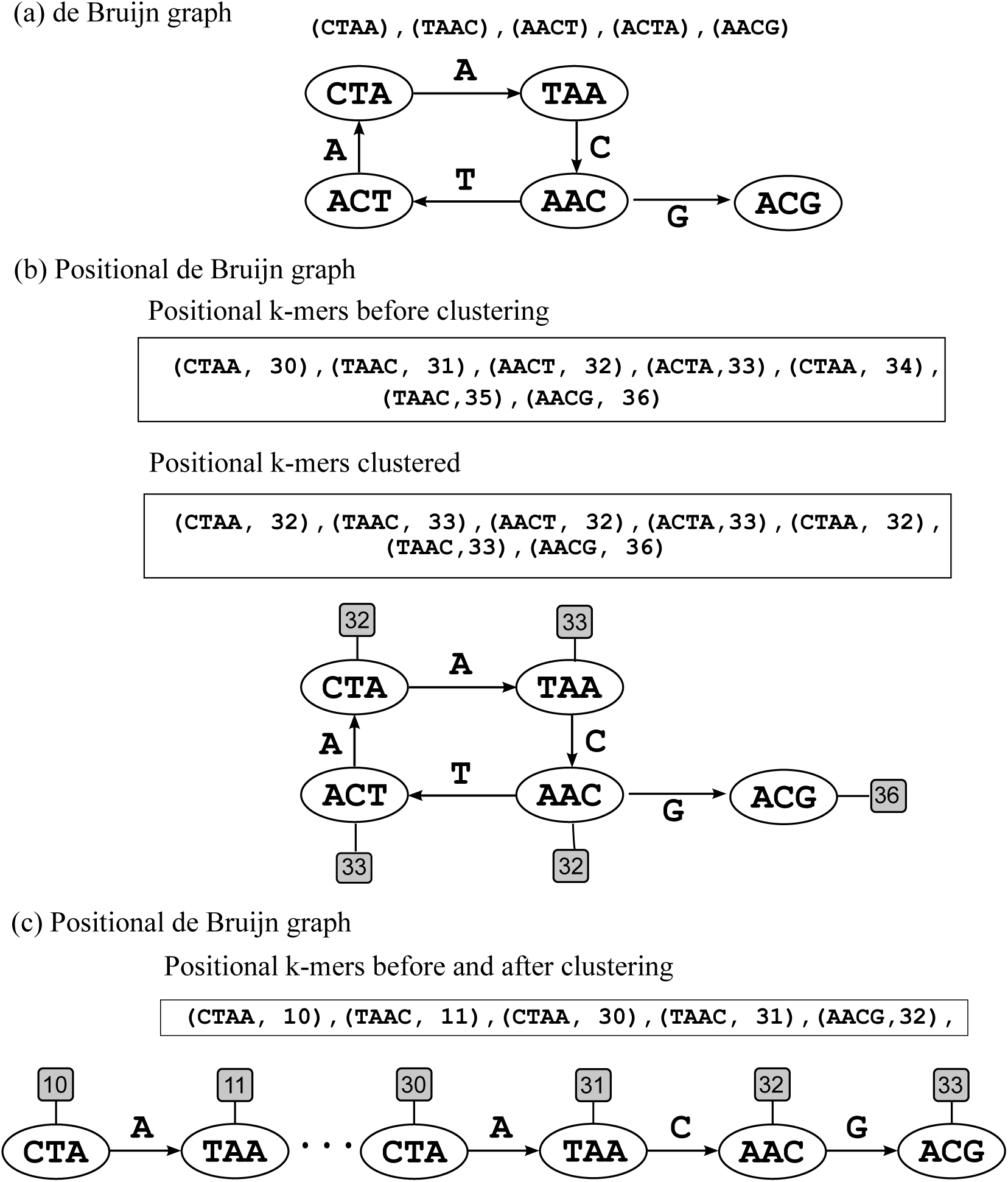
An illustration showing when a whirl in the positional de Bruijn graph can prevail. (a) shows a de Bruijn graph constructed for a read CTAACTAACG and *k* = 4. (b) shows the positional de Bruijn graph with constructed for a read CTAACTAACG whose alignment starts at position 30 of the genome, *k* = 4 and *Δ* = 4. The set of positional *k*-mers before and after clustering with Δ are illustrated. (c) shows the positional de Bruijn graph with constructed for a read CTAA..CTAACG whose alignment starts and resumes at position 10 and 30 of the genome respectively. In this last example, *k* = 4 and *Δ* = 4. As can be seen by these illustrations, whirls will persist in the positional de Bruijn graph for short genomic repeats when the difference between *k* and *Δ* is reasonably small since they will create positional *k*-mers whose multiplicity is greater than one. In (b) CTAA at position 32 and TAAC at position 33 both have multiplicity 2 after clustering.

The existence of whirls in the positional de Bruijn graph is possible, but extremely rare. Figure 3 illustrates when a whirl can persist in a positional de Bruijn graph. In this example, (a) and (b) illustrate the de Bruijn graph and the positional de Bruijn graph constructed for *k* = 4 and *Δ* = 4 and read CTAACTAACG that aligns to position 30 in the genome. Both the de Bruijn graph and its positional counterpart contain a whirl. The whirl in the positional de Bruijn graph is created since the occurrences of CTAA and TAAC are clustered together at positions 32 and 33, respectively, creating positional *k*-mers that have multiplicity greater than one. The third graph in Figure 3 shows, more typically, how positional information resolves the whirls within the graph.

Our main contribution has been to demonstrate the effect of adding positional information to long read assembly and how optical mapping data can assist in the assembly of long reads. Given the rarity of whirls in the positional de Bruijn graph, we expect that even slightly more sophisticated traversal algorithms would be capable of constructing 94% or more of the *E. coli* genome with only a few contigs that have a small mismatch rate (Koota has 1.71 mismatches per 100 kbp) without using more than 1.2GB of space. This would bridge the gap between long-read and short-read assembly since it would enable longer (more complicated genomes) to be assembled the same accuracy as short reads.

A further advantage of integrating the positional information into the de Bruijn graph is that it allows for a meaningful partitioning of the graph. Each partition of the graph would contain the *k*-mers belonging to an interval of positions. Each of these partitions could be independently processed yielding a natural way to develop parallel or distributed algorithms for the positional de Bruijn graph.

## Footnotes

1 OpGen (Correspondence:http://www.opgen.com) and BioNano (http://www.bionanogenomics.com) are commercial producers of optical mapping data.

2 https://github.com/P50acificBiosciences/DevNet/wiki/Saccharomyces-cerevisiae-W303-Assembly-Contigs

## References

1. A. Bankevich, S. Nurk, D. Antipov, A.A. Gurevich, M. Dvorkin, A.S. Kulikov, V.M. Lesin, S.I. Nikolenko, S. Pham, A. D. Prjibelski, A.V. Pyshkin, A. V. Sirotkin, N. Vyahhi, G. Tesler, M.A. Alekseyev, and P. A. Pevzner. SPAdes: A new genome assembly algorithm and its applications to single-cell sequencing. Journal of Computational Biology, 19(5):455–477, 2012.

2. K. Berlin, S. Koren, C.-S. Chin, J.P. Drake, J.M. Landolin, and A.M. Phillippy. Assembling large genomes with single-molecule sequencing and locality-sensitive hashing. Nature Biotechnology, 33:623–630, 2015.

3. A. Bowe, T. Onodera, K. Sadakane, and T. Shibuya. Succinct de Bruijn graphs. In Proc. WABI, pages 225–235, 2012.

4. M. Burrows and D.J. Wheeler. A block sorting lossless data compression algorithm. Technical Report 124, Digital Equipment Corporation, 1994.

5. R. Chikhi, A. Limasset, S. Jackman, J.T. Simpson, and P. Medvedev. On the representation of de Bruijn graphs. In Proc. RECOMB, pages 35–55, 2014.

6. R. Chikhi and G. Rizk. Space-efficient and exact de Bruijn graph representation based on a Bloom filter. Algorithms for Molecular Biology, 8(22), 2012.

7. C.-S. Chin, D.H. Alexander, P. Marks, A.A. Klammer, J. Drake, C. Heiner, A. Clum, A. Copeland, J. Huddleston, E.E. Eichler, S.W. Turner, and J. Korlach. Nonhybrid, finished microbial genome assemblies from long-read smrt sequencing data. Nature Methods, 10(6):563–569, 2013.

8. T.C. Conway and A.J. Bromage. Succinct data structures for assembling large genomes. Bioinfomatics, 27(4):479–486, 2011.

9. Y. Dong, M. Xie, Y. Jiang, N. Xiao, X. Du, W. Zhang, G. Tosser-Klopp, J. Wang, S. Yang, J. Liang, W. Chen, J. Chen, P. Zeng, Y. Hou, C. Bian, S. Pan, Y. Li, X. Liu, W. Wang, B. Servin, B. Sayre, B. Zhu, D. Sweeney, R. Moore, W. Nie, Y. Shen, R. Zhao, G. Zhang, J. Li, T. Faraut, J. Womack, Y. Zhang, J. Kijas, N. Cockett, X. Xu, S. Zhao, J. Wang, and W. Wang. Sequencing and automated whole-genome optical mapping of the genome of a domestic goat *(Capra hircus)*. Nature Biotechnology, 31(2):135–141, 2013.

10. S. Gog, T. Beller, A. Moffat, and M. Petri. From theory to practice: Plug and play with succinct data structures. In Proc. SEA, pages 326–337, 2014.

11. A. Gurevich, V. Saveliev, N. Vyahhi, and G. Tesler. QUAST: Quality assessment tool for genome assemblies. Bioinformatics, 29(8):1072–1075, 2013.

12. R.M. Idury and M.S. Waterman. A new algorithm for DNA sequence assembly. Journal of Computational Biology, 2:291–306, 1995.

13. S. Koren and A.M. Phillippy. One chromosome, one contig: complete microbial genomes from long-read sequencing and assembly. Current Opinion in Microbiology, 23:110–120, 2015.

14. S. Koren, B.P. Walenz, K. Berlin, J.R. Miller, and A.M. Phillippy. Canu: scalable and accurate long-read assembly via adaptive k-mer weighting and repeat separation. bioRxiv, 2016.

15. A.K.-Y. Leung, T.-P. Kwok, R. Wan, M. Xiao, P.-Y. Kwok, K.Y. Yip, and T.-F. Chan. OMBlast: Alignment tool for optical mapping using a seed-and-extend approach. Bioinformatics, 2016. To appear.

16. Y. Lin, M.W. Shen, J. Yuan, M. Chaisson, and P.A. Pevzner. Assembly of long error-prone reads using de bruijn graphs. In Proc. RECOMB, page 265, 2016.

17. L.M. Mendelowitz, D.C. Schwartz, and M. Pop. MAligner: a fast ordered restriction map aligner. Bioinformatics, 32(7):1016–1022, 2016.

18. E.W. Myers, G.G. Sutton, A.L. Delcher, et al. A whole-genome assembly of drosophila. Science, 287:2196–2204, 2000.

19. N. Nagarajan, T.D. Read, and M. Pop. Scaffolding and validation of bacterial genome assemblies using optical restriction maps. Bioinformatics, 24(10):1229–1235, 2008.

20. D. Okanohara and K. Sadakane. Practical entropy-compressed rank/select dictionary. In In Proc. ALENEX, pages 60–70, 2007.

21. Y. Ono, K. Asai, and M. Hamada. PBSIM: PacBio reads simulator—toward accurate genome assembly. Bioinformatics, 29(1):119–121, 2013.

22. M. Pendleton, R. Sebra, A.W.C. Pang, et al. Assembly and diploid architecture of an individual human genome via single-molecule technologies. Nature Methods, 12:780–786, 2015.

23. P.A. Pevzner, H. Tang, and G. Tesler. De novo repeat classification and fragment assembly. Genome Research, 14(9):1786–1796, 2004.

24. P.A. Pevzner, H. Tang, and M.S. Waterman. An Eulerian path approach to DNA fragment assembly. Proceedings of the National Academy of Sciences of the United States of America, 98(17):9748–9753, 2001.

25. A. Rhoads and K.F. Au. PacBio sequencing and its applications. Genomics, Proteomics & Bioinformatics, 13(5):278–289, 2015.

26. L. Salmela, R. Walve, E. Rivals, and E. Ukkonen. Accurate self-correction of errors in long reads using de Bruijn graphs. Bioinformatics, 2016. To appear.

27. J.T. Simpson and R. Durbin. Efficient construction of an assembly string graph using the FM-index. Bioinformatics, 26(12):i367–i373, 2010.

28. J.T. Simpson, K. Wong, S.D. Jackman, J.E. Schein, S.J.M. Jones, and I. Birol. ABySS: A parallel assembler for short read sequence data. Genome Research, 19(6):1117–1123, 2009.

29. A. Valouev, L. Li, Y.-C. Liu, D.C. Schwartz, Y. Yang, Y. Zhang, and M.S. Waterman. Alignment of optical maps. Journal of Computational Biology, 13(2):442–462, 2006.

30. D. Verzotto, A.S.M. Teo, A.M. Hillmer, and N. Nagarajan. OPTIMA: Sensitive and accurate whole-genome alignment of error-prone genomic maps by combinatorial indexing and technology-agnostic statistical analysis. GigaScience, 5:2, 2016.

31. D.R. Zerbino and E. Birney. Velvet: Algorithms for de novo short read assembly using de Bruijn graphs. Genome Research, 18(5):821–829, 2008.

